# Iris Morphological and Biomechanical Factors Influencing Angle Closure During Pupil Dilation

**DOI:** 10.1101/2024.04.24.591028

**Authors:** Royston K.Y. Tan, Gim Yew Ng, Tin A. Tun, Fabian A. Braeu, Monisha E. Nongpiur, Tin Aung, Michael J.A. Girard

## Abstract

**Purpose:** To use finite element (FE) analysis to assess what morphological and biomechanical factors of the iris and of the anterior chamber are more likely to influence angle narrowing during pupil dilation.

**Methods:** The study consisted of 1,344 FE models comprising of the cornea, sclera, lens and iris (stroma, sphincter and dilator tissues) to simulate pupil dilation and to assess changes in angle. For each model, we varied the following parameters: anterior chamber depth (ACD = 2 –4 mm) and width (ACW = 10–12 mm), iris convexity (IC = 0–0.3 mm), thickness (IT = 0.3–0.5 mm), stiffness (*E* = 4–24 kPa) and Poisson’s ratio (*v* = 0–0.3), and simulated pupil dilation. We evaluated for the change in anterior chamber angle (△∠) and the final dilated anterior chamber angles (∠_f_) from baseline to dilation for each parameter.

**Results:** The final dilated AC angles decreased with a smaller ACD (∠_f_ = 53.4°±12.3° to 21.3°±14.9°), smaller ACW (∠_f_ = 48.2°±13.5° to 26.2°±18.2°), larger IT (∠_f_ = 52.6°±12.3° to 24.4°±15.1°), larger IC (∠_f_ = 45.0°±19.2° to 33.9°±16.5°), larger *E* (∠_f_ = 40.3°±17.3° to 37.4°±19.2°) and larger *v* (∠_f_ = 42.7°±17.7° to 34.2°±18.1°).

The change in AC angle increased with larger ACD (△∠ = 9.37°±11.1° to 15.4°±9.3°), smaller ACW (△∠ = 7.4°±6.8° to 16.4°±11.5°), larger IT (△∠ = 5.3°±7.1° to 19.3°±10.2°), smaller IC (△∠ = 5.4°±8.2° to 19.5°±10.2°), larger *E* (△∠ = 10.9°±12.2° to 13.1°±8.8°) and larger *v* (△∠ = 8.1°±9.4° to 16.6°±10.4°).

**Conclusions:** This parametric study offered valuable insights into the factors that could influence angle closure. The morphology of the iris (IT and IC) and its innate biomechanical behavior (*E* and *v*) were crucial in influencing the way the iris deformed during dilation, and angle closure was further exacerbated by decreased AC biometry (ACD and ACW).

## Introduction

Angle closure is characterized by the closure of the anterior chamber (AC) angle between the iris and the cornea, blocking aqueous outflow through the trabecular meshwork.^1, 2^ This may occur due to mechanisms such as pupillary block and plateau iris, or conditions that affect the anatomy such as cataracts or ciliary body tumors. Angle narrowing occurs especially when the pupil dilates, pushing the iris towards the periphery and blocking the AC angle. The factors associated with primary angle closure glaucoma (PACG) have been extensively explored through imaging techniques such as optical coherence tomography^3^ (OCT) and ultrasound biomicroscopy^4^ (UBM) to determine the significance of each anterior segment biometric parameter. Amongst these are several morphological parameters associated with ACG:^1, 5, 6^ shallow AC depth (ACD), smaller AC width (ACW), larger iris thickness (IT) (i.e., IT500, IT750, IT2000 and ITM), thicker lens vault, and shorter axial length But given that angle closure is a very dynamic process, it is also plausible to suspect that the biomechanics of the iris would play a significant role.

However, thus far, no studies have investigated the iris morphological and biomechanical factors influencing angle closure during pupil dilation. The iris is a highly robust tissue, with constant fluid exchange through its pores^7, 8^ when responding to nerve inputs, and the knowledge of its biomechanical behavior may help us understand why angle closure occurs. An approach to depicting this alteration in volume involves using Poisson’s ratio,^9^ where a higher ratio indicates reduced fluid mobility and a potential association with angle closure. It is our belief that angle closure occurrence is contributed by both morphological and biomechanical factors. In fact, past studies had shown differences in iris biomechanical properties between normal and PACG patients,^10^ including increased stiffness and decreased permeability.^3^ These observations could be explained by alterations in densities of collagen type I and III^11, 12^ which would increase the tissue stiffness, and an increase in extracellular matrix would in turn decrease the porosity of the iris tissue.

To conduct a thorough investigation into the impact of iris biomechanics, it would be necessary to capture images of a vast and varied cohort of individuals undergoing pupil dilation. Accomplishing this task in a real-world setting presents significant difficulties. Nonetheless, the advent of digital twin technology offers a viable alternative to study an extensive range of hypothetical patient profiles.^13^ Specifically, in this research, our aim was to employ finite element (FE) analysis to determine which morphological and biomechanical attributes of the iris and AC are most important in inducing angle closure when the pupil dilates.

## Methods

A total of 1,344 FE models were evaluated for the AC and angles before and after pupil dilation with varying anterior chamber parameters. Each finite element model was designed as a quadrant of an axisymmetric AC across two planes, with the assumption of a perfectly spherical eye. The model was first constructed using SolidWorks and exported into Abaqus for meshing. Finally, the meshed geometry was imported into FEBio for analysis in all scenarios. For this study, we varied 4 AC parameters and 2 iris biomechanical parameters for analysis: ACD, ACW, IT, iris convexity (IC), iris stiffness (*E*) and iris Poisson’s ratio (*v*).

## 3D Model

The AC model consisted of the cornea, sclera, lens, iris (consisting of the stroma, sphincter muscle and dilator muscle) and ciliary body. The model, created with SolidWorks (2022, Dassault Systèmes, Vélizy-Villacoublay, France), was designed based on literature values: cornea thickness of 0.50 mm,^14, 15^ cornea radius of curvature of 7.50 mm,^16^ sclera radius of curvature of 12.00 mm^17^ and only the anterior lens of radii of 1.4 mm and 5.00 mm (**Figure 1A and 1B**). The cornea and sclera were combined as a single tissue since the stiffness of the tissues was similar and far greater than the iris (essentially a rigid body). The trabecular meshwork region was also combined with the cornea and sclera, and angled 15° from the cornea at a small curvature (**Figure 1B and 1C**). The lens, intended to be a rigid body for iris sliding during pupil dilation, was simplified as a shell with a thickness of 0.10 mm and an inner diameter of 2.00 mm.

**Figure 1.**
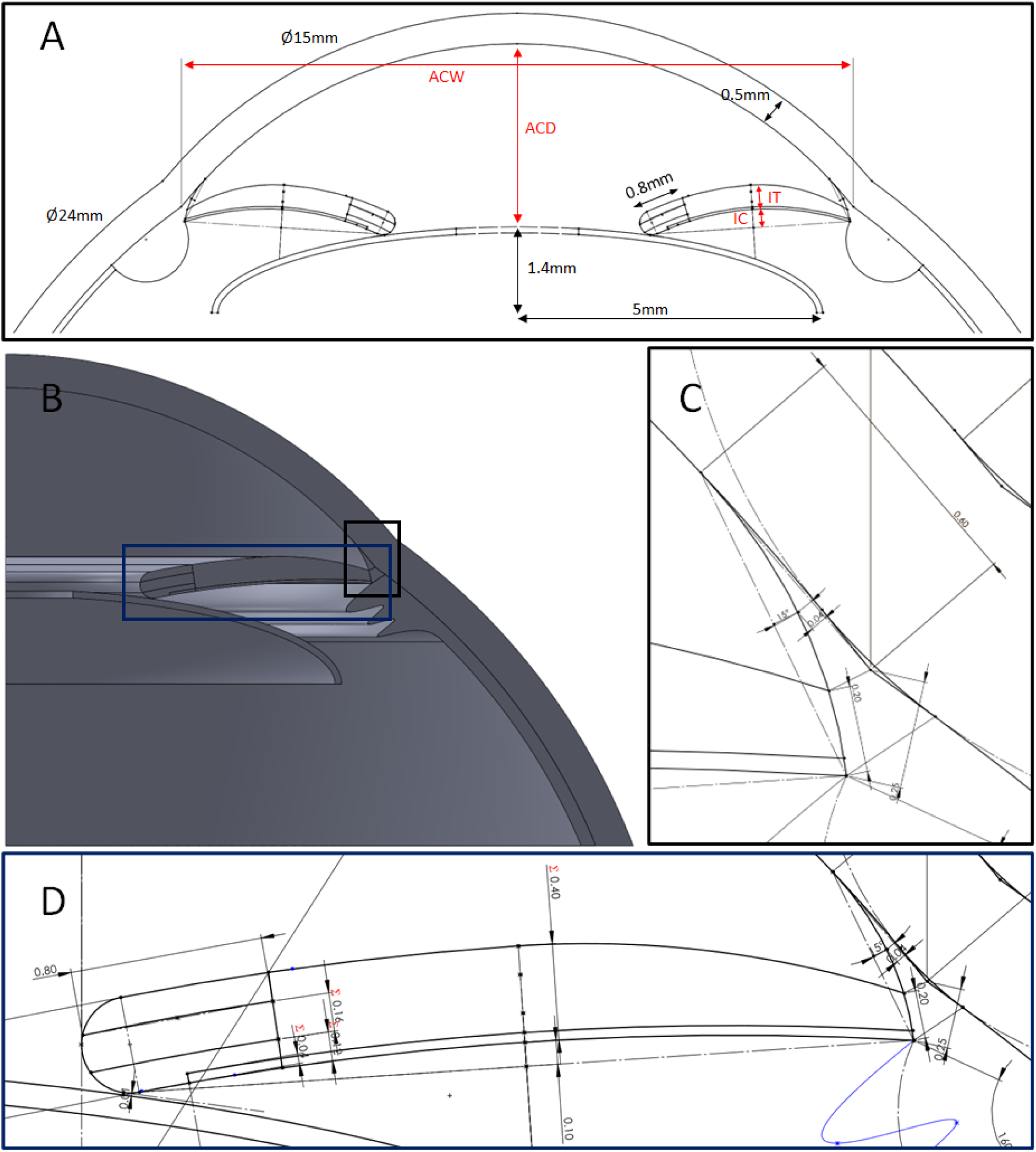
(**A**) Anterior chamber parameters for the models, with the analyzed parameters highlighted in red. **(B)** The CAD model of the anterior chamber was designed to align with anatomical measurements. **(C)** Simplification of the model by combining the trabecular meshwork, cornea and scleral regions since they did not affect the final results. **(D)** The iris was designed with a 0.20 mm root, sphincter muscle at two-fifths of the thickness and 0.80 mm radius and the dilator muscle at one-tenth the iris thickness beginning from the mid-radius of the sphincter muscle to the iris root.

The iris was attached to the sclera at the root, which was kept consistent for all cases at 0.20 mm. Iris tissue thickness was defined as the thickness from the iris margin to mid-periphery, and it decreased from the mid-periphery to the iris root. The iris tissue was rounded at the margin and consisted of the sphincter muscle at two-fifths of the thickness and a 0.80 mm radial length. The dilator muscle was located at the posterior of the iris, with a thickness one-tenth that of the iris. The dilator muscle spanned from the mid-portion of the sphincter muscle to the iris root (**Figure 1D**). A ciliary body was included for illustration purposes and did not influence the simulation results.

### Mesh and Convergence

The 3D model was exported from SolidWorks as a STEP file (AP203) and imported into Abaqus FEA (2021, Dassault Systèmes, Vélizy-Villacoublay, France) for meshing. Eight-node hexahedral elements were predominantly used for each model, unless otherwise stated.

A convergence study was performed on an average model with mesh sizes of 6,800, 23,730, 53,120 and 344,960, and convergence test showed that the selected mesh density was < 0.8% of the most refined mesh results and was deemed numerically acceptable.

### Biomechanical Properties

The finite element model contained 6 parts with the following material parameters: The corneo-scleral shell was described with an incompressible neo-Hookean formulation with a Young’s modulus of 500 kPa.^18^ The lens was described as a rigid body, and fixed for all scenarios. The sphincter and dilator muscles were described as incompressible neo-Hookean materials with a Young’s modulus of 40 kPa assigned to both. The dilator muscle was represented using the solid mixture material model comprised of a prescribed active uniaxial muscle contraction force and a passive hyperelastic modulus. The AC angles were extracted when the iris was dilated by 1.1 mm for all cases. The direction of contraction was in the iris radial direction, which was specified using the element local coordinates. Finally, the iris stroma was also described as a neo-Hookean material, with varying biomechanical properties according to our sensitivity study described below. Of note, the neo-Hookean formulation was selected primarily for its stability and accuracy when performing large deformations in FEBio.

### Boundary Conditions

Constraints along the axisymmetric planes of the quadrant AC model were enforced, and a zero-friction contact interface was assumed between the iris and lens, as well as between the iris and the corneal endothelium surface. The lens was fixed, and so was the outer boundary of the corneo-scleral shell.

### Varying the Parameters – FE Sensitivity Study

According to population studies and estimates, the anterior chamber biometry can vary considerably, even amongst healthy individuals. This study aimed to understand the extent of influence of AC parameters in angle closure. To do so, we used a design of experiments^19^ approach to investigate how the variations in ACD, ACW, IT, IC, *E* and *v* could influence the final AC angle and change in AC angle during pupil dilation. ACD was defined as the distance between the cornea endothelium to the lens, IT was defined as the thickness at mid-periphery, and IC was defined by the perpendicular distance of the line joining the iris base and the iris tip at midpoint (**Figure 1A**). Instead of the horizontal distance between the scleral spurs, ACW was defined as the distance between the points where the cornea and sclera radii intersected (**Figure 1A**). Each parameter was assigned either 3 or 4 values using the design of experiments method including the lower and upper limits: ACD = 2, 3 & 4 mm,^20–22^ ACW = 10, 11 & 12 mm,^21, 23^ IT = 0.3, 0.4 & 0.5 mm,^24–26^ IC = 0, 0.1, 0.2 & 0.3 mm,^24, 27^ *E* = 4, 8, 14 & 24 kPa^3, 28^ and *v* = 0, 0.1, 0.2 & 0.3.^3, 29, 30^

### FE Processing to Predict Anterior Chamber Angles

All FE models were solved using FEBio (FEBio Studio v1.8.2, University of Utah, UT, USA), a non-linear FE solver designed for biomechanical studies. The AC angles were determined using the coordinates of the nodes at the scleral spur and its adjacent nodes to calculate the initial and final angles, and we reported the change in AC angles (Δ∠) as well as the final AC angles (∠_f_). For each parameter, Δ∠ and ∠_f_ were reported as the average and standard deviation of all the scenarios with the parameter value, and the *p*-values were determined using the two-sample t-Test assuming unequal variances. The cases were classified based on the final AC angle: open angles had final AC angles of ∠_f_ > 20°, narrow angles had final AC angles of 0° < ∠_f_ < 20°,^23, 31^ and angle closure cases had final AC angles of ∠_f_ = 0°. Multiple linear regression analysis was performed to rank the effects of each evaluated parameter. We normalized each parameter from 0 – 1 and reported the results as the coefficients ± standard errors. The multiple linear regression was governed by the equation:

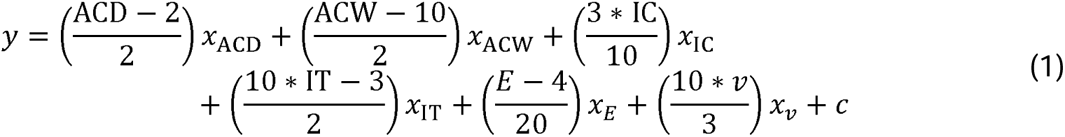

where *y* is the independent variable (Δ∠, ∠_f_ or percentage Δ∠), *x* is the coefficient of the dependent parameter and *c* is the constant. Since the variance of the parameters were skewed for angle closure cases (∠_f_ ≥ 0°), we performed statistical analysis using the Wilcoxon rank sum test and employed the Bonferroni correction.

## Results

### Classification and Angle Changes

Of the 1,344 simulated cases of pupil dilation, 61 (4.54%) ended up with angle closure, 160 (11.9%) with narrow angles and 1,123 (83.56%) maintained open angles. During pupil dilation, the AC angle width decreased for all scenarios, but the magnitude of change varied based on the parameter, without influencing the final AC angle (**Figures 2 and 3**). When ACD decreased from 4 mm to 2 mm, the ∠_f_ decreased following pupil dilation (53.4° ± 12.3° to 21.3° ± 14.9°; *P* < 0.001), even though Δ∠ decreased (15.4° ± 9.3° to 9.37° ± 11.1°; *P* < 0.001). Similarly, when ACW decreased from 12 mm to 10 mm, the ∠_f_ decreased following pupil dilation (48.2° ± 13.5° to 26.2° ± 18.2°; *P* < 0.001), and Δ∠ increased (7.4° ± 6.8° to 16.4° ± 11.5°; *P* < 0.001). The largest changes were seen when IT increased from 0.3 mm to 0.5 mm, where ∠_f_ decreased the most (52.6° ± 12.3° to 24.4° ± 15.1°; *P* < 0.001), and Δ∠ increased (5.3° ± 7.1° to 19.3° ± 10.2°; *P* < 0.001). When IC increased from 0 mm to 0.3 mm, the ∠_f_ decreased following pupil dilation (45.0° ± 19.2° to 33.9° ± 16.5°; *P* < 0.001), even though Δ∠ decreased (19.5° ± 10.2° to 5.4° ± 8.2°; *P* < 0.001). The smallest changes were seen in stroma stiffness *E*, when increased from 4 kPa to 24 kPa, the ∠_f_ decreased following pupil dilation (40.3° ± 17.3° to 37.4° ± 19.2°; *P* = 0.0644), and Δ∠ increased (10.9° ± 12.2° to 13.1° ± 8.8°; *P* < 0.001). When Poisson’s ratio *v* increased from 0 to 0.3, the ∠_f_ decreased following pupil dilation (42.7° ± 17.7° to 34.2° ± 18.1°; *P* < 0.001), and Δ∠ increased (8.1° ± 9.4° to 16.6° ± 10.4°; *P* < 0.001).

**Figure 2.**
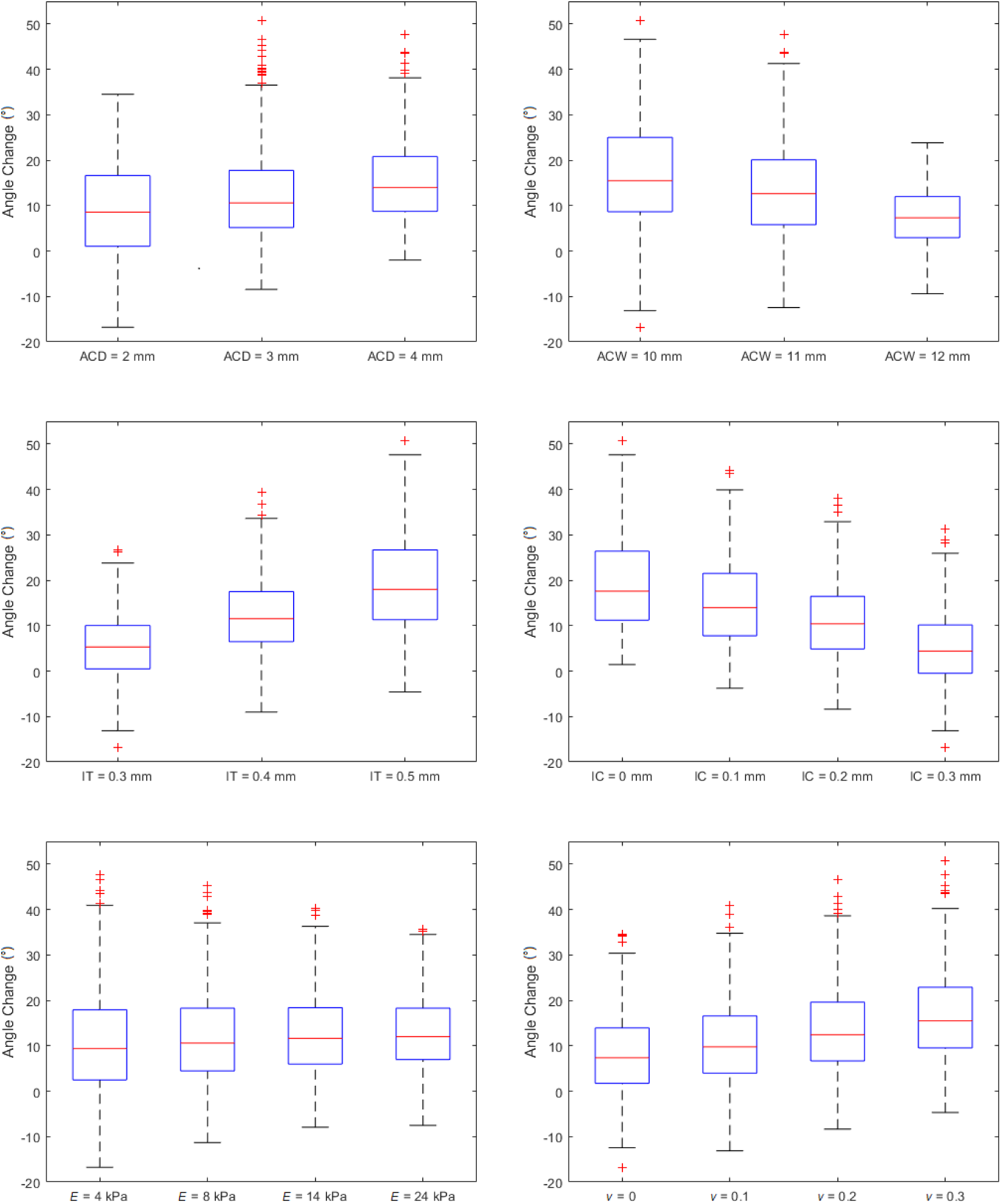
Box plots of the angle changes with respect to the explored parameters. Note that the positive angle changes represent a decrease in the anterior chamber angle. There were statistical intergroup differences for the parameters ACD, ACW, IT, IC and *v*, and *E* = 4 kPa and 24 kPa (*P* < 0.05).

**Figure 3.**
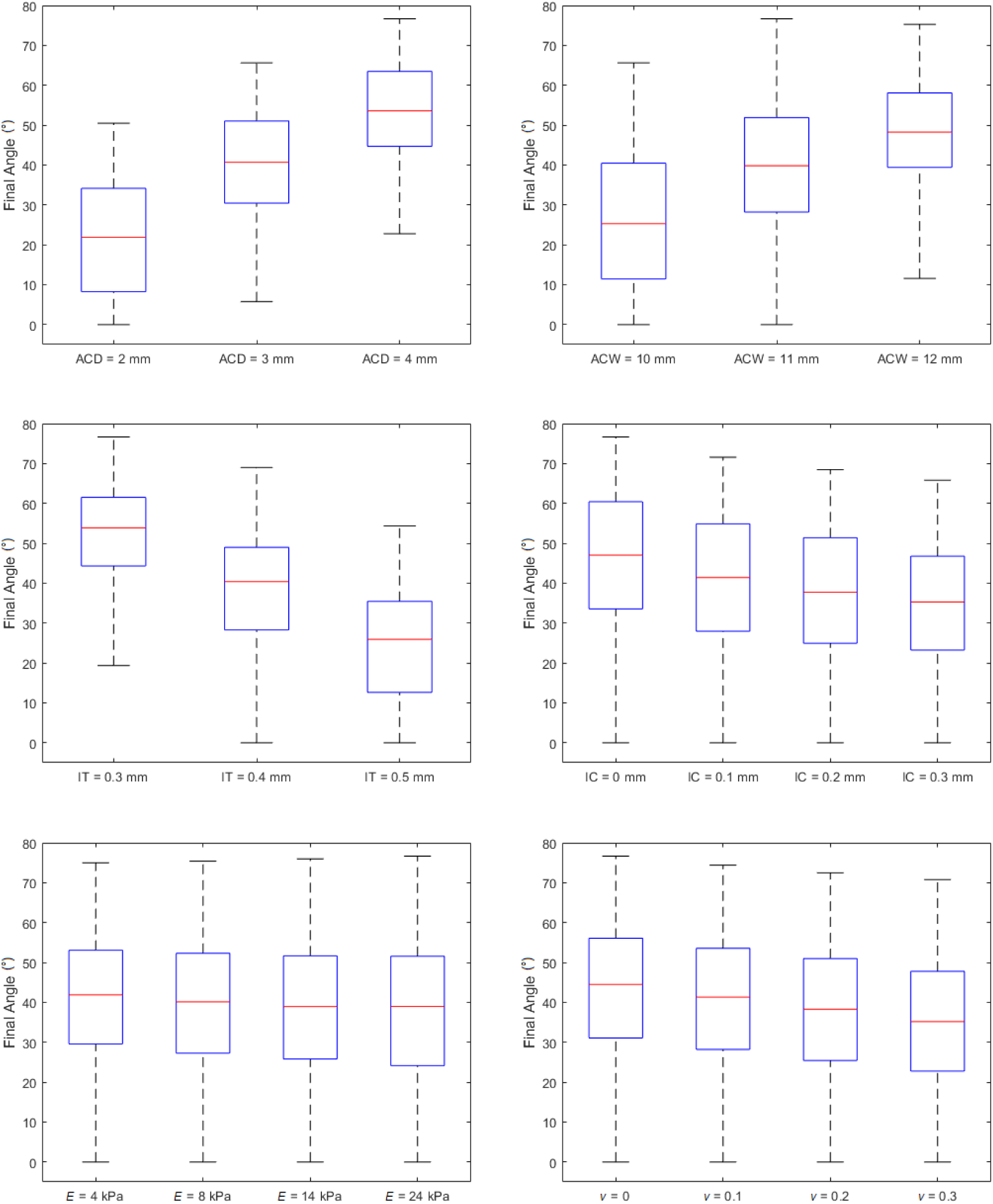
Box plots of the final anterior chamber angles with respect to the explored parameters. There were statistical intergroup differences for the parameters ACD, ACW, IT, IC and *v* (*P* < 0.05).

### Multiple Linear Regression

The results of the multiple linear regression showed that, for change in AC angles (the AC angles always decrease after pupil dilation), the order of influence of the parameter coefficients were as follows: IT (14.0 ± 0.269, *P* < 0.001), ACW (−12.6 ± 0.315, *P* < 0.001), IC (−12.0 ± 0.301, *P* < 0.001), ACD (10.0 ± 0.326, *P* < 0.001), *v* (8.51 ± 0.295, *P* < 0.001) and *E* (1.90 ± 0.292, *P* < 0.001). The constant was 4.98 ± 0.347 and the adjusted R^2^ was 0.85. Note that the negative values indicate an inverse relationship (e.g., increase in IT results in larger decrease in AC angles, but increase in ACW results in lesser decrease in AC angles) (**Figure 4A**).

**Figure 4.**
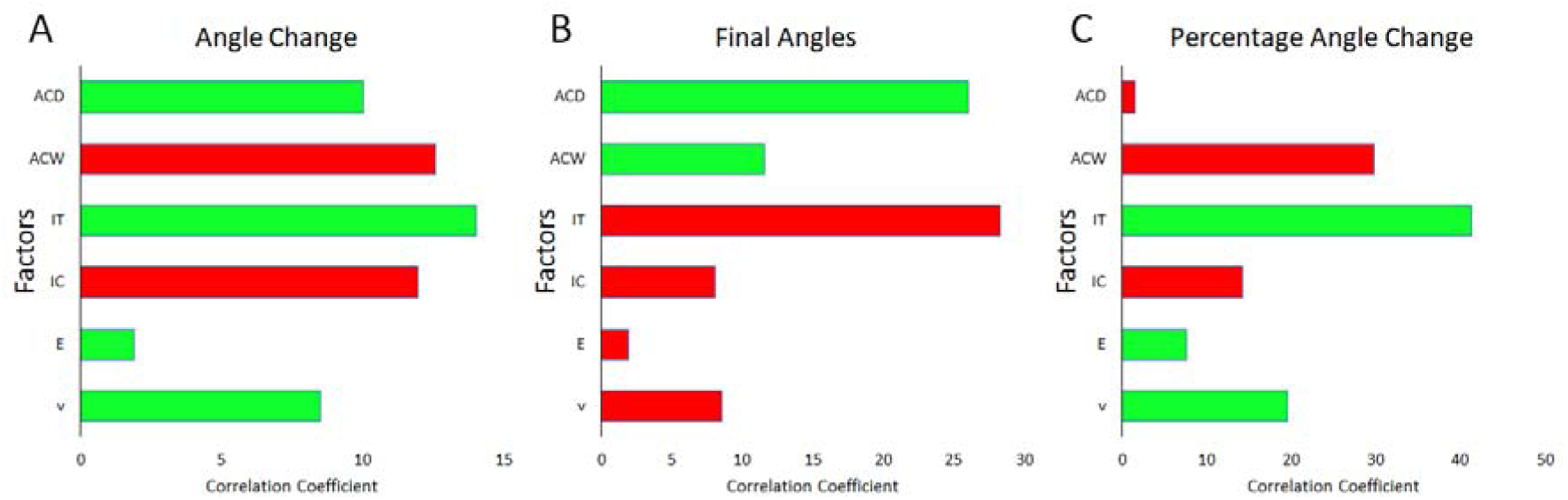
Multiple linear regression of AC Parameters with respect to (**A**) Angle change, (**B**) Final angles and (**C**) Percentage change from the initial angle. The bars in green represent a positive correlation with the factor, while the bars in red represent a negative correlation with the factor.

For the final AC angles, the order of influence of the parameter coefficients were as follows: IT (−28.2 ± 0.287, *P* < 0.001), ACD (26.0 ± 0.347, *P* < 0.001), ACW (11.6 ± 0.335, *P* < 0.001), *v* (−8.51 ± 0.314, *P* < 0.001), IC (−8.06 ± 0.321, *P* < 0.001) and *E* (−1.90 ± 0.311, *P* < 0.001). The constant was 45.9 ± 0.370 and the adjusted R^2^ was 0.944. Note that the negative values indicate an inverse relationship (e.g., increase in IT results in decrease in final AC angles) (**Figure 4B**).

For the percentage change in angles (with respect to the initial angle), the order of influence of the parameter coefficients are as follows: IT (41.3 ± 0.978, *P* < 0.001), ACW (− 29.8 ± 1.14, *P* < 0.001), *v* (19.5 ± 1.07, *P* < 0.001), IC (−14.2 ± 1.09, *P* < 0.001), *E* (7.62 ± 1.06, *P* < 0.001) and ACD (−1.58 ± 1.18, *P* = 0.734). The constant was 9.44 ± 1.26 and the adjusted R^2^ was 0.71 Note that the negative values indicate an inverse relationship (e.g., increase in ACW results in a smaller percentage change in angle) (**Figure 4C**).

## Discussion

Our study combined clinical knowledge with computational analysis to understand how AC dynamics could influence angle closure. Our models were able to predict cases of angle closure, revealing that the AC angle width always decreased during pupil dilation. The extent of this angle change varied based on the combination of parameters that had changed, with specific parameters such as iris thickness having greater impact.

### Increased Iris Thickness, Convexity, Stroma Stiffness and Poisson’s Ratio Lead to Narrower AC Angles

Existing literature was only able to report significance for individual factors that correlate with angle closure, and our FE predictions were able to highlight the disproportionate influence of the 6 investigated parameters in relation to the AC angle. It was clear from the results that the morphology of the iris (thickness and convexity) and its innate biomechanical behavior (stiffness and Poisson’s ratio) were crucial in influencing the way the iris deformed during dilation, and angle closure was further exacerbated by decreased AC biometry (ACD and ACW).

Our results showed that iris thickness had the largest effect on the narrowing of the chamber angle. This result corroborated literature findings, specifically reports of iris thickness at 750 µm, 2000 µm and maximum (IT750, IT2000 and ITM)^24, 25^ from the sclera spur. As the primary tissue involved in angle closure, it is unsurprising that iris morphology is directly related to narrower AC angles.

Iris convexity had also been correlated with angle closure, with increased convexity reported in patients with ACG.^24, 32^ This phenomenon is likely to be the result of muscle activity controlling pupil size, the biomechanical differences between the stroma and muscle, as well as the aqueous pressure that is induced between the anterior and posterior chambers when the dilator muscle contract.^33^ Hence, iris convexity is influenced by the biomechanical equilibrium between the stroma and muscle states within the AC, and our results also indicated that the stiffness and Poisson’s ratio of the stroma were correlated to angle closure.

### Our FE Model Was Able to Predict Cases of Angle Closure

In our study, we had designed the AC to be accurate and robust, to have few geometrical compromises and able to perform the high deformation simulations (**Figure 5**). Our results showed that a small fraction (4.54%) of the cases had angle closure. Among these 61 scenarios, all of them had a shallow AC (ACD = 2 mm), 59 had a thick iris (IT = 0.5 mm) and 58 had a narrow AC width (ACW = 10 mm). It is important to note that this percentage of cases with angle closure applies to the prescribed dilator muscle contraction force dilating the pupil from 4.0 mm to about 5.1 mm, and we expect the number of scenarios for angle closure to increase when the iris is dilated further. Our FE presents an opportunity to develop a tool for angle closure, and we believe that clinical diagnosis could be improved by combining the knowledge from clinical imaging techniques and computational biomechanics to evaluate for angle closure. Using tools such as inverse FE,^3^ it may be possible to improve prognosis of angle closure, allowing potential candidates to take precautionary measures for the disease.

**Figure 5.**
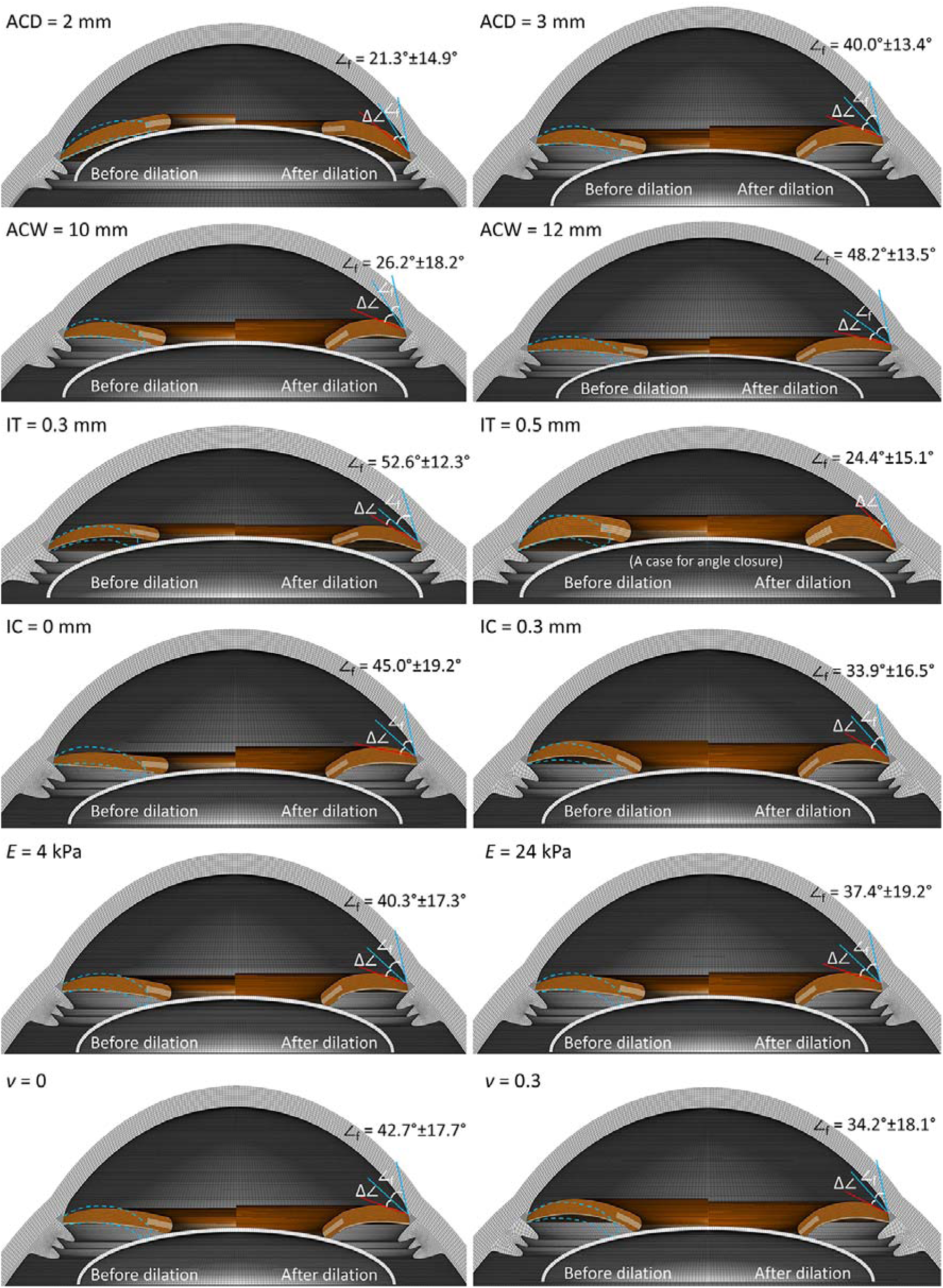
Comparing AC parameters with significant correlation: ACD, ACW, IT, IC, *E* and *v*. For each AC figure, the left and right halves show the iris before and after dilator muscle contraction. The red and blue lines show the initial and final AC angles, and the dotted blue lines show the outline of the final iris shape. ∠_f_ is presented as the (mean ± SD) of cases corresponding to the parameter.

### The Amount of Angle Change Does Not Reflect the Final AC Angle

Whether angle closure develops depends on the interaction with many parameters, and it is clear from the multiple linear regression coefficients that the correlations are not straightforward. For example, when IT increases from 0.3 mm to 0.5 mm, the Δ∠ increases and the ∠_f_ decreases. However, when ACD increases from 2 mm to 4 mm, the Δ∠ increases and ∠_f_ increases as well. This meant that a thicker iris contributed more to closing of the AC angle, yet a larger AC angle from a larger ACD was effective in resisting angle closure caused by the other factors. Therefore, the state of the final AC angle is dependent on the combination of all the factors, with the extreme cases (i.e., large IT, shallow ACD, small ACW and high *v*) resulting in closure of the chamber angle.

### Anterior Chamber Interactions is Highly Complex

There are several more parameters highlighted in literature which were found to have strong correlations with angle closure.^34–37^ Two of such parameters are the anterior chamber area (ACA), and by geometric transitive property (i.e., 2D to 3D), the anterior chamber volume (ACV). These are dependent parameters from the cumulative effect of other AC biometry (i.e., cornea radius of curvature, sclera radius of curvature, ACD, ACW, IT, IC, lens thickness, etc.) influencing the residual ACA/ACV space. Likewise, the combined effect of iris biomechanical properties (i.e., *E* and *v*) could vary the ACA/ACV changes during pupil dilation. In fact, Foo et al.^38^ ranked determinants influencing AC angles and found ACA/ACV to be most significantly correlated, along with lens vault (LV), which is also influenced by AC biometry (i.e., ACD, ACW, lens thickness, etc.). This implied that the dynamic interactions within the AC are immensely complex and likely to be a cumulative effect of all factors at disproportionate weightage, with inter-dependencies of parameters affecting one another. We believe more work is required in segregating and analyzing individual parameters to correlate them with angle closure.

Additionally, whilst we analyze 2D images captured, it is important to remember that there are many concurrent processes happening, and the equilibrium state of the tissues within the AC can be difficult to ascertain. At the exterior, extraocular muscles are constantly changing the position of the globe, and saccadic eye movements create fluctuations of cellular stresses^39^ from the inertia of the aqueous humor, which in turn affects iris movements. During distance accommodation, ciliary muscle contraction pushes the ciliary body centrally and anteriorly to increase lens curvature and increases the lens vault.^40–43^ This in turn pushes the iris anteriorly during near vision. During light accommodation, the iris dilator muscle constricts when light intensity decreases to increase pupil size, pushing the anterior border layer closer to the cornea. All these accommodative processes not only changes the positions of the intraocular tissues, but also the differential pressure between the anterior and posterior chambers of the anterior segment of the eye.^33^ The invention of laser peripheral iridotomy seeks to prevent angle closure by relieving pupil block and excessive iris anterior bowing.^44^ These anterior chamber interactions are highly complex and dynamic, and whilst we attempt to rationalize individual measurable parameters, it is important to remember these parameters often influence one another, and this parametric study represents the first step in gaining a clearer understanding of the development of angle closure.

### Limitations

As with all studies, the assumptions and design choices of this study had some drawbacks. First, we performed this study using the ‘design of experiments’ method. This approach allowed us to systematically distribute cases among all the parameters. However, while this method provides a valuable framework for understanding the influence of each parameter, it does not reflect the distribution of these parameters in a human population, which is likely to follow a normal or skewed distribution.^45^ This implies that the design of experiments method could introduce higher result variance, potentially reducing the probability values in correlations that might have been significant. With a normal or skewed distribution, we also expect the cases of angle closure to be greatly reduced, since the prevalence of these extreme cases is at the tailed end of the population distribution.

Second, our computational study examined the solid mechanics of soft tissues in the AC. The conditions *in vivo* could be affected by aqueous humor production, its dynamic interactions with the iris and outflow through the trabecular meshwork and ciliary body (conventional and unconventional pathways)^46^. The difference in pressure between the anterior and posterior chambers had been hypothesized to influence iris convexity and AC angles,^27^ but it could prove to be challenging to model such biphasic simulations with solid-fluid interfaces. Additionally, the aforementioned biometry parameters may influence iris dynamics, including lens curvature, crypts and furrows that would interact with the aqueous humor and affect how the pupil dilates.

Third, research on iris biomechanics is lacking, especially at the microscopic level. The iris anterior border layer (ABL) has been investigated to reveal an inter-crimp fiber pattern, with the fiber direction approximately 45 degrees away from the radial direction. Below the ABL, the sparse collagen network is much less understood, with little information regarding the porosity and biomechanical properties of the stroma meshwork. This makes the model sensitive to inaccuracies in material properties, which could have an impact on the results.

Finally, we generated models based on general geometric shapes, each with a smooth ABL and uniform iris curvature. The models were not able to account for variable features such as non-uniform iris thickness and curvature, iris crypts and furrows. These features affect iris dilation movement and explain cases of short iridotrabecular contact cases^47^ and concave iris cases,^48^ further contributing to the incidence of angle closure.

## Conclusions

This study highlights the intricate role of AC biometry and biomechanics influencing angle closure, underscoring the importance of multi-parameter analysis for the development of more accurate diagnostic techniques. While our study utilized ‘hypothetical’ patients, combining this approach with optical coherence tomography imaging in a clinical scenario could enhance the identification of patients at risk of developing angle closure. Further research is essential to enhance the complexity of models, ultimately improving clinical prognosis for preventing the development of angle closure cases.

---

**Table 1.** Change in AC angle and final AC angles w.r.t parameter changes. The positive change in angle denotes a decrease of the AC angle.

| Parameter | | Change in AC angle ($\Delta\angle$) | Final AC angle ($\angle_f$) |
| --- | --- | --- | --- |
| ACD (mm) | 2 | $9.4^\circ \pm 11.1^\circ$ | $21.3^\circ \pm 14.9^\circ$ |
| | 3 | $12.2^\circ \pm 10.1^\circ$ | $40.0^\circ \pm 13.4^\circ$ |
| | 4 | $15.4^\circ \pm 9.3^\circ$ | $53.4^\circ \pm 12.3^\circ$ |
| ACW (mm) | 10 | $16.4^\circ \pm 11.5^\circ$ | $26.2^\circ \pm 18.2^\circ$ |
| | 11 | $13.2^\circ \pm 10.5^\circ$ | $39.6^\circ \pm 16.3^\circ$ |
| | 12 | $7.4^\circ \pm 6.8^\circ$ | $48.2^\circ \pm 13.5^\circ$ |
| IT (mm) | 0.3 | $5.3^\circ \pm 7.1^\circ$ | $52.6^\circ \pm 12.3^\circ$ |
| | 0.4 | $12.2^\circ \pm 8.5^\circ$ | $38.6^\circ \pm 14.8^\circ$ |
| | 0.5 | $19.3^\circ \pm 10.2^\circ$ | $24.4^\circ \pm 15.1^\circ$ |
| IC (mm) | 0 | $19.5^\circ \pm 10.2^\circ$ | $45.0^\circ \pm 19.2^\circ$ |
| | 0.1 | $15.1^\circ \pm 9.4^\circ$ | $39.8^\circ \pm 18.3^\circ$ |
| | 0.2 | $11.1^\circ \pm 8.5^\circ$ | $36.9^\circ \pm 17.4^\circ$ |
| | 0.3 | $5.4^\circ \pm 8.2^\circ$ | $33.9^\circ \pm 16.5^\circ$ |
| $E$ (kPa) | 4 | $10.9^\circ \pm 12.2^\circ$ | $40.3^\circ \pm 17.3^\circ$ |
| | 8 | $12.3^\circ \pm 10.7^\circ$ | $38.6^\circ \pm 18.1^\circ$ |
| | 14 | $12.8^\circ \pm 9.6^\circ$ | $37.9^\circ \pm 18.7^\circ$ |
| | 24 | $13.0^\circ \pm 8.8^\circ$ | $37.4^\circ \pm 19.2^\circ$ |
| $\nu$ | 0 | $8.1^\circ \pm 9.4^\circ$ | $42.7^\circ \pm 17.7^\circ$ |
| | 0.1 | $10.8^\circ \pm 9.8^\circ$ | $40.0^\circ \pm 18.0^\circ$ |
| | 0.2 | $13.6^\circ \pm 10.1^\circ$ | $37.1^\circ \pm 18.1^\circ$ |
| | 0.3 | $16.6^\circ \pm 10.4^\circ$ | $34.2^\circ \pm 18.1^\circ$ |

## References

1. Nongpiur ME, Ku JY, Aung T. Angle closure glaucoma: a mechanistic review. Current opinion in ophthalmology 2011;22:96–101.

2. Quigley HA, Friedman DS, Congdon NG. Possible mechanisms of primary angle-closure and malignant glaucoma. Journal of glaucoma 2003;12:167–180.

3. Panda SK, Tan RK, Tun TA, et al. Changes in iris stiffness and permeability in primary angle closure glaucoma. Investigative Ophthalmology & Visual Science 2021;62:29–29.

4. Pavlin CJ, Foster FS. Ultrasound biomicroscopy of the eye: Springer Science & Business Media; 2012.

5. Wright C, Tawfik MA, Waisbourd M, Katz LJ. Primary angle-closure glaucoma: an update. Acta ophthalmologica 2016;94:217–225.

6. Amerasinghe N, Aung T. Angle-closure: risk factors, diagnosis and treatment. Progress in brain research 2008;173:31–45.

7. Quigley HA. The iris is a sponge: a cause of angle closure. Ophthalmology 2010;117:1–2.

8. Freddo TF. Ultrastructure of the iris. Microscopy research and technique 1996;33:369–389.

9. Pant AD, Dorairaj SK, Amini R. Appropriate objective functions for quantifying iris mechanical properties using inverse finite element modeling. Journal of Biomechanical Engineering 2018;140:074502.

10. Pant AD, Gogte P, Pathak-Ray V, Dorairaj SK, Amini R. Increased iris stiffness in patients with a history of angle-closure glaucoma: an image-based inverse modeling analysis. Investigative ophthalmology & visual science 2018;59:4134–4142.

11. Konstas A, Marshall G, Lee W. Immunocytochemical localisation of collagens (I–V) in the human iris. Graefe’s archive for clinical and experimental ophthalmology 1990;228:180–186.

12. Chung C, Dai M, Lin J, Wang Z, Chen H, Huang J. Correlation of iris collagen and in-vivo anterior segment structures in patients in different stages of chronic primary angle-closure in both eyes. Indian Journal of Ophthalmology 2019;67:1638.

13. Hollingsworth K, Bowyer KW, Flynn PJ. Pupil dilation degrades iris biometric performance. Computer vision and image understanding 2009;113:150–157.

14. Sadoughi MM, Einollahi B, Einollahi N, Rezaei J, Roshandel D, Feizi S. Measurement of central corneal thickness using ultrasound pachymetry and Orbscan II in normal eyes. Journal of ophthalmic & vision research 2015;10:4.

15. Bovelle R, Kaufman SC, Thompson HW, Hamano H. Corneal thickness measurements with the Topcon SP-2000P specular microscope and an ultrasound pachymeter. Archives of Ophthalmology 1999;117:868–870.

16. Khng C, Osher RH. Evaluation of the relationship between corneal diameter and lens diameter. Journal of Cataract & Refractive Surgery 2008;34:475–479.

17. Augusteyn RC, Nankivil D, Mohamed A, Maceo B, Pierre F, Parel J-M. Human ocular biometry. Experimental eye research 2012;102:70–75.

18. Whitford C, Studer H, Boote C, Meek KM, Elsheikh A. Biomechanical model of the human cornea: considering shear stiffness and regional variation of collagen anisotropy and density. Journal of the mechanical behavior of biomedical materials 2015;42:76–87.

19. Atkinson AC, Hunter WG. The design of experiments for parameter estimation. Technometrics 1968;10:271–289.

20. Yuen LH, He M, Aung T, Htoon HM, Tan DT, Mehta JS. Biometry of the cornea and anterior chamber in Chinese eyes: an anterior segment optical coherence tomography study. Investigative ophthalmology & visual science 2010;51:3433–3440.

21. Leung C, Palmiero P, Weinreb R, et al. Comparisons of anterior segment biometry between Chinese and Caucasians using anterior segment optical coherence tomography. British journal of ophthalmology 2010;94:1184–1189.

22. Jonas JB, Nangia V, Gupta R, et al. Anterior chamber depth and its associations with ocular and general parameters in adults. Clinical & experimental ophthalmology 2012;40:550–556.

23. Lee RY, Chon BH, Lin S-C, He M, Lin SC. Association of ocular conditions with narrow angles in different ethnicities. American journal of ophthalmology 2015;160:506–515. e501.

24. Wang B, Sakata LM, Friedman DS, et al. Quantitative iris parameters and association with narrow angles. Ophthalmology 2010;117:11–17.

25. Wang B, Narayanaswamy A, Amerasinghe N, et al. Increased iris thickness and association with primary angle closure glaucoma. British Journal of Ophthalmology 2011;95:46–50.

26. He M, Wang D, Console JW, Zhang J, Zheng Y, Huang W. Distribution and heritability of iris thickness and pupil size in Chinese: the Guangzhou Twin Eye Study. Investigative ophthalmology & visual science 2009;50:1593–1597.

27. Nonaka A, Iwawaki T, Kikuchi M, Fujihara M, Nishida A, Kurimoto Y. Quantitative evaluation of iris convexity in primary angle closure. American journal of ophthalmology 2007;143:695–697.

28. Whitcomb JE, Barnett VA, Olsen TW, Barocas VH. Ex vivo porcine iris stiffening due to drug stimulation. Experimental eye research 2009;89:456–461.

29. Mak H, Xu G, Leung CK-S. Imaging the iris with swept-source optical coherence tomography: relationship between iris volume and primary angle closure. Ophthalmology 2013;120:2517–2524.

30. Tun TA, Baskaran M, Perera SA, et al. Sectoral variations of iridocorneal angle width and iris volume in Chinese Singaporeans: a swept-source optical coherence tomography study. Graefes Arch Clin Exp Ophthalmol 2014;252:1127–1132.

31. Huang G, Gonzalez E, Peng P-H, et al. Anterior chamber depth, iridocorneal angle width, and intraocular pressure changes after phacoemulsification: narrow vs open iridocorneal angles. Archives of ophthalmology 2011;129:1283–1290.

32. Da Soh Z, Thakur S, Majithia S, Nongpiur ME, Cheng C-Y. Iris and its relevance to angle closure disease: a review. British Journal of Ophthalmology 2020.

33. Silver DM, Quigley HA. Aqueous flow through the iris–lens channel: Estimates of differential pressure between the anterior and posterior chambers. Journal of glaucoma 2004;13:100–107.

34. Guzman CP, Gong T, Nongpiur ME, et al. Anterior segment optical coherence tomography parameters in subtypes of primary angle closure. Investigative ophthalmology & visual science 2013;54:5281–5286.

35. Nongpiur M, Atalay E, Gong T, et al. Anterior segment imaging-based subdivision of subjects with primary angle-closure glaucoma. Eye 2017;31:572–577.

36. Chansangpetch S, Rojanapongpun P, Lin SC. Anterior segment imaging for angle closure. American journal of ophthalmology 2018;188:xvi–xxix.

37. Wu R-Y, Nongpiur ME, He M-G, et al. Association of narrow angles with anterior chamber area and volume measured with anterior-segment optical coherence tomography. Archives of ophthalmology 2011;129:569–574.

38. Foo L-L, Nongpiur ME, Allen JC, et al. Determinants of angle width in Chinese Singaporeans. Ophthalmology 2012;119:278–282.

39. Abouali O, Modareszadeh A, Ghaffarieh A, Tu J. Investigation of saccadic eye movement effects on the fluid dynamic in the anterior chamber. Journal of biomechanical engineering 2012;134:021002.

40. Findl O, Kiss B, Petternel V, et al. Intraocular lens movement caused by ciliary muscle contraction. Journal of Cataract & Refractive Surgery 2003;29:669–676.

41. Koke MP. Mechanism of accommodation. Archives of Ophthalmology 1942;27:950–968.

42. Fernández-Vigo JI, Kudsieh B, Shi H, De-Pablo-Gómez-de-Liaño L, Fernández-Vigo JÁ, García-Feijóo J. Diagnostic imaging of the ciliary body: Technologies, outcomes, and future perspectives. European Journal of Ophthalmology 2022;32:75–88.

43. Drexler W, Baumgartner A, Findl O, Hitzenberger CK, Fercher AF. Biometric investigation of changes in the anterior eye segment during accommodation. Vision Research 1997;37:2789–2800.

44. He M, Jiang Y, Huang S, et al. Laser peripheral iridotomy for the prevention of angle closure: a single-centre, randomised controlled trial. The Lancet 2019;393:1609–1618.

45. Chen H, Lin H, Lin Z, Chen J, Chen W. Distribution of axial length, anterior chamber depth, and corneal curvature in an aged population in South China. BMC ophthalmology 2016;16:1–7.

46. Johnson M, McLaren JW, Overby DR. Unconventional aqueous humor outflow: a review. Experimental eye research 2017;158:94–111.

47. Mizoguchi T, Ozaki M, Wakiyama H, Ogino N. Peripheral iris thickness and association with iridotrabecular contact after laser peripheral iridotomy in patients with primary angle-closure and primary angle-closure glaucoma. Clinical ophthalmology 2014;517–522.

48. Cheung CY-l, Liu S, Weinreb RN, et al. Dynamic analysis of iris configuration with anterior segment optical coherence tomography. Investigative ophthalmology & visual science 2010;51:4040–4046.

